# Perturbed cyclic strain results in a mechanically-induced, premature senescent phenotype in cardiac fibroblasts

**DOI:** 10.1101/2025.03.16.643588

**Authors:** Stephanie E. Schneider, Adrienne K. Scott, Katie M. Gallagher, Emily Y. Miller, Soham Ghosh, Corey P. Neu

**Affiliations:** Paul M. Rady Department of Mechanical Engineering, University of Colorado Boulder, CO; Biomedical Engineering Program, University of Colorado Boulder, CO; Department of Mechanical Engineering, Colorado State University, Ft. Collins, CO, USA; School of Biomedical Engineering, Colorado State University, Ft. Collins, CO, USA; Translational Medicine Institute, Colorado State University, Ft. Collins, CO, USA; BioFrontiers Institute, University of Colorado Boulder, CO

## Abstract

The cardiovascular system operates under continuous cyclic mechanical stretch, and changes in mechanics and biochemical responses observed in cardiovascular disorders activate the cardiac fibroblasts (CFs) and increase cellular senescence in the tissue. However, it is unclear if cellular senescence is initiated from biomechanical stimulus alone. Here, we subjected murine CFs to uniaxial stretch and perturbed the mechanical stimulation to examine the impact on induction of a senescent phenotype. Loss of stretch magnitude and increase in frequency, mimicking an injurious hypertrophic or fibrotic response, led to a senescence-like phenotype, including cell cycle, lamin B expression, and DNA damage. Mechanical induction of CF senescence relied on p53/p21, whereas CF senescence induction triggered by reactive oxygen species or involving a mutation in lamin A/C gene occurred through p16. Moreover, mechanical induction of premature senescence was associated with decreases in the nuclear envelope protein emerin. These results demonstrate perturbed mechanical stimulation can initiate a senescent state and altered nuclear integrity may initiate this phenotype.

**Teaser:** Perturbed mechanics initiates a premature senescent-like phenotype in cardiac fibroblasts.

## Introduction

Cellular senescence is a state of cell cycle arrest that plays a role in tissue formation and remodeling during development and following injury (*1*, *2*). It can lead to reduced tissue regeneration and function which contributes to inflammation and promote tumorigenesis in aging organisms. While traditionally characterized as an arrest in cell cycle, the hallmarks of senescence have been more broadly defined which include DNA damage response, secretory associated senescent phenotype, and apoptosis resistance (*3–5*). Extrinsic factors such as oncogene disruption, irradiation, chemotherapy, developmental cues, and tissue damage have been shown to generate different forms of stress-induced senescence (*1*). To further complicate the identification of a senescent phenotype, no one single biomarker is specific for every induction of the senescent pathway (*6*). Instead, multiple biochemical signals, morphological features, and alterations in protein and gene expression are used to define the senescent phenotype.

The most extensively used markers to define senescence are the p16/p19 in mice (or p16/p14 in humans) and p53/p21 pathways for cell cycle arrest (*4*, *7*). The p53/21 axis is also activated with the DNA damage response signaling pathway which can give rise to the senescent phenotype. Biophysical features such as the enlargement of the cell and nuclear area, and increased granularity within the cell occur in addition to biochemical signaling and gene expression changes. Interestingly, many of the hallmarks associated with the senescent phenotype overlap with biophysical alterations associated with the nucleus such as global changes in histone modifications and chromatin remodeling, and changes at the nuclear envelope such as the loss of lamin B within the cell (*8*, *9*). Finally, increased excretion of inflammatory cytokines, chemokines, and matrix metalloproteinases generate a persistent feedback loop which further directs the cellular pathway from a return to the homeostatic state or toward apoptosis (*2*). Due to the numerous pathways, heterogeneous senescent phenotypes, and the discovery of new initiators of senescence, the characterization of this cellular state needs to be defined in a tissue and disease-dependent manner (*10*).

Emerging evidence has shown altered mechanical stimulus and microenvironments could provide another factor to initiate a premature senescent-like state (*11*, *12*). With prolonged exposure to physical stresses or stretch deformation as observed in trauma and disease, the cell and nucleus undergo direct changes that influence an array of biomarkers (*13–16*). However, if the cellular stress continues outside of a normative physiological range (*17*), a premature stress-induced senescent phenotype can emerge. This phenotype is linked with disease states such as cancer, osteoarthritis, and cardiac disease (*3*, *18*). Notably, disease states often lead to changes in both biochemical and mechanical environments in the tissue indicating senescence may initiate from a combination of factors.

Due to the many disorders associated with cardiovascular diseases such as hypertension, cardiomyopathy, heart failure, and cardiac hypertrophy, this disease state has become the leading cause of death worldwide (*19*, *20*). One common symptom of cardiovascular diseases is fibrosis which arises from excessive deposition of extracellular matrix proteins (*21*, *22*). The stiffer fibrotic tissue affects cellular contractions in a complex manner which in turn leads to decreased tissue stretch and increased frequency of contraction, and therefore, directly impacts the nucleus (*15*, *23*–*25*). Cardiac fibroblasts (CFs) are one of the main cell types involved in the adaptive response to environmental cues in heart tissue. CFs regulate the remodeling and repair process, and the balance between healthy or fibrotic tissue (*26*). Recent studies have shown the phenotypic plasticity of the CF through mechanosensitive responses both biochemical through calcium signaling and biophysically through increased α-SMA (*27–29*). Additionally, alterations in stretch magnitude and stiffness of the microenvironment are known to activate cardiac fibroblasts into a myofibroblast phenotype (*30*, *31*). These studies highlight the numerous mechanisms driving CF activation and underscore the relevance of better understanding the stress responses of the CF, such as cellular senescence, in the context of cardiovascular diseases (*32*). Cellular senescence and elimination of senescent cells have been examined as a potential therapy in cardiovascular diseases (*21*, *33*, *34*). Senescent cardiomyocytes display contractile dysfunction due to reduced cellular contractions which results in an increased contractile frequency (*21*). Nested between the cardiomyocytes, the cardiac fibroblasts passively experience this contractile dysfunction, this suggests a continuous crosstalk between the cardiomyocyte and cardiac fibroblasts (*35*). Moreover, senescent CFs have been observed in cardiac tissue several days after myocardial infarction which led us to hypothesize about the mode of senescent induction in CFs especially after both a mechanically and biochemically-stressed event (*36*, *37*).

To study how contractile dysfunction influences the CF phenotype, we use a reductionist approach to mimic the mechanical disruption described in cardiac disease using an *in vitro* model. Specifically, we examined CFs if perturbed mechanical stimulation can induce a premature stress-induced phenotype in primary murine CFs by examining established markers of cellular senescence. We further investigated the senescent response of CFs through oxidative stress and CFs with a lamin A/C mutation as alterations in structural integrity of the nuclear envelope and increased oxidative stress is associated with several types of cardiovascular disease (*38–40*). Analysis of the mechanoresponsive nature of the CF revealed that a premature senescent phenotype can be initiated from perturbations in the mechanical environment alone and is unique when compared to chemical induction.

## Results

### Perturbed mechanical stimulation initiates a senescent-like phenotype

To examine the role that perturbed mechanical stimulation has on cardiac fibroblasts (CFs), we plated primary murine wild type (WT) cardiac fibroblasts on flexible membranes and subjected them to two modes of cyclic stretch for 4 days (**Fig. 1A**). The first mode, termed **Stretch Control**, was 8.5% strain at 1 Hz with daily medium changes. The second mode, termed **Stretch Injury**, which mimicked altered mechanical loading observed in cardiac disease (*30*, *31*) was 24 hours at 8.5% strain at 1 Hz, and then the strain was decreased to 2.5% strain and frequency increased to 2 Hz with daily medium changes (**Fig. 1B**). The decreased strain and increased frequency loading mode were selected to mimic the contractile dysfunction observed in cardiac disease in both humans and mice. After 4 days (96 hours), stretched cells were analyzed with gene expression and immunofluorescence for markers of cell cycle arrest and alterations in the nuclear envelope. When comparing Stretch Control CFs to Stretch Injury CFs, we observed a decrease in proliferation. Specifically, using Ki-67 that labels cells in G1, G2, and S phase of the cell cycle, Stretch Injury CFs showed a significant decrease in the Ki-67 normalized intensity values compared to Stretch Control CFs (**Fig. 1C**), this suggesting a generic senescence phenotype caused by Stretch Injury.

**Figure 1:**
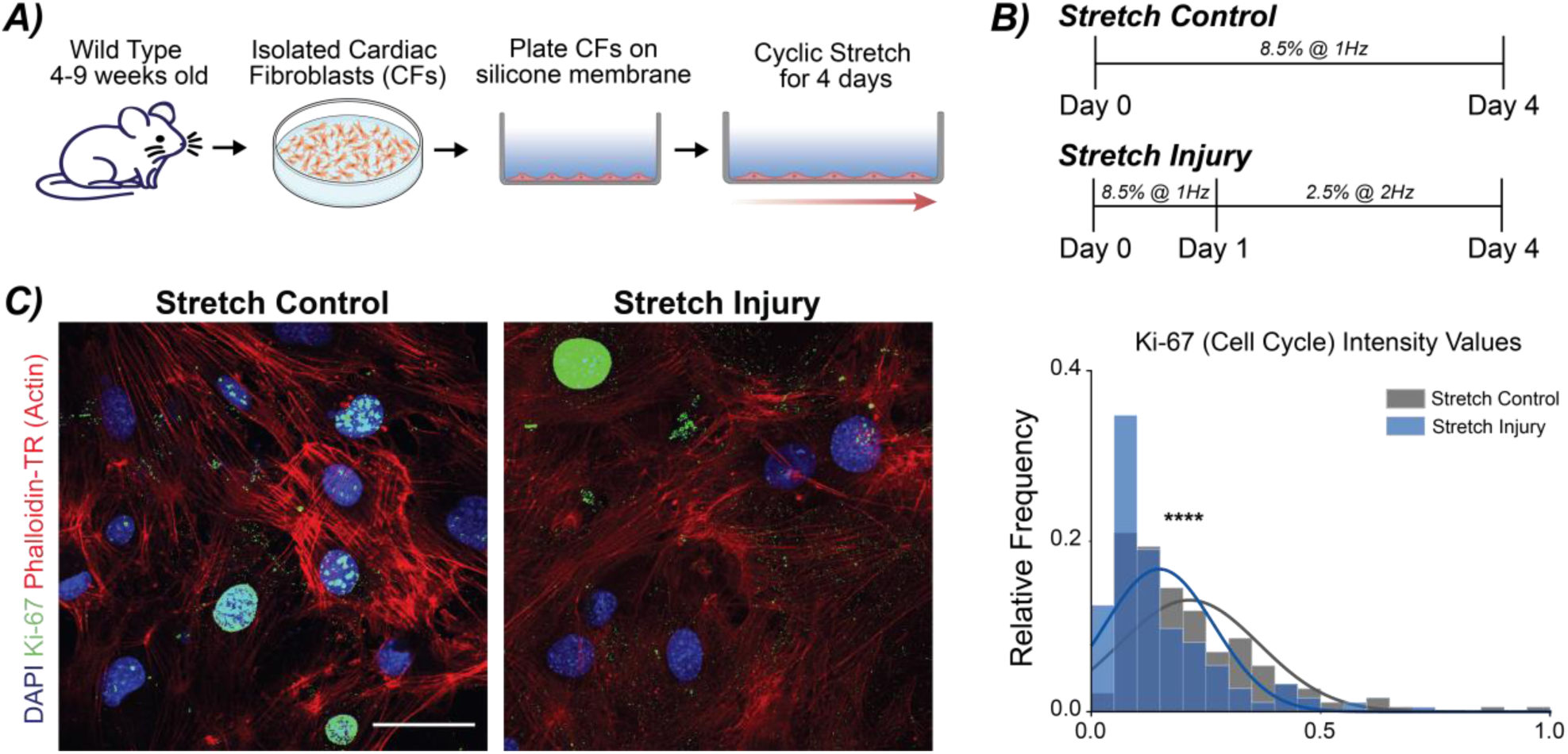
Perturbed loading disrupts the cell cycle and proliferation in primary cardiac fibroblasts. (A) Primary murine cardiac fibroblasts (CFs) were isolated from wild type mice between 4-9 weeks. Cells were expanded on TCP before plating on Geltrex-coated silicone membranes. CFs were cyclically stretched for 4 days with two separate loading schemes. (B) Two loading schemes were used to mimicking the transition in mechanical stretch and frequency changes observed in cardiovascular disease. (C) Representative images show Ki-67 staining in Stretch Control and Stretch Injury cultures. The histogram depicts the relative frequency of Ki-67 staining between the two loading schemes. A decrease in Ki-67 positive cells were observed with Stretch Injury, noting a decrease in proliferation. DAPI = blue (405nm), Ki-67 = green (488nm), Actin (Phalloidin-TR) = red (561nm). Scale bar = 50µm. N = 5 animals, n ≥ 25 nuclei/treatment. ****p <0.0001.

Next, we assessed Stretch Injury CFs for other hallmarks of senescence such as lamin B expression, alterations in nuclear morphology, DNA damage, and gene expression for markers of cell cycle arrest. Decreased lamin B is observed in senescent cells both *in vivo* and *in vitro* (*41*). To evaluate lamin B levels through immunofluorescence, we quantified the thickness of the lamin B ring as a surrogate measure of lamin B protein expression. Stretch Injury CFs had a significantly decreased lamin B ring compared Stretch Control CFs (**Fig. 2A**). Nuclear area significantly increased in Stretch Injury CFs (**Fig. 2B**). However, nuclear aspect ratio changes were not observed (**fig. S1A**). Surprisingly, Stretch CFs had increased chromatin condensation and H3K9me3 foci even though other studies have shown chromatin relaxation and decreases in H3K9me3 with senescent phenotypes (*42*) (**fig. S1B** and **C**). Next, we examined the Stretch Injury CFs for DNA damage through staining for ɣH2A.X which is associated with increased double-strand breaks (DSB) of the DNA and localization of DNA damage response markers are frequently observed in premature-stress induced senescence (*5*). Using a custom MATLAB code, we quantified the number of ɣH2A.X foci per nucleus for each stretch mode. By measuring DNA damage through the presence of ɣH2A.X which localizes to DSB in the nucleus, we found a significant increase in ɣH2A.X foci per nucleus in the Stretch Injury CFs compared to the Stretch Control CFs (**Fig. 2C**). The increased ɣH2A.X in Stretch Injury CFs could provide one signaling mechanism that triggers the initiation of the senescent state in the CFs.

**Figure 2:**
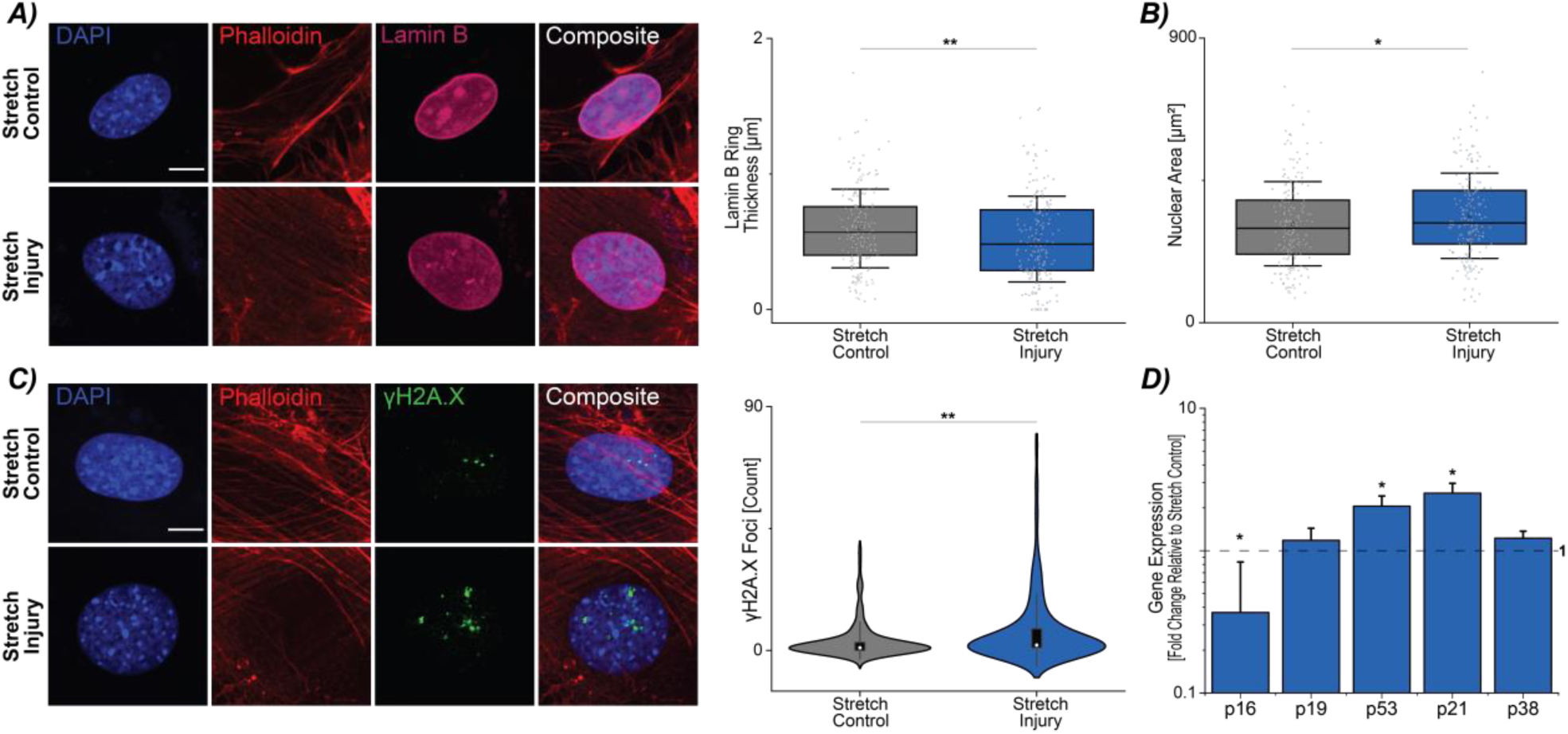
Cardiac fibroblasts experiencing perturbed loading display markers indicative of a mechanically-induced senescent phenotype, including alterations in morphology, DNA damage, and gene expression. (A) Cyclically stretched CFs in the control and injury loading schemes were stained for lamin B. Expression of lamin B was assessed by quantifying the ring thickness around the nucleus with stretch injury cultures having a decrease compared to stretch controls. Scale bar = 10µm. Error bar = 1 Std. (B) Stretch injury CFs had a decrease in nuclear area compared to stretch controls. Error bar = 1 Std. (C) γH2A.X foci were counted per nucleus between the two loading schemes. Stretch injury CFs had increased foci compared to stretch controls. Error bar = 1 Std. Scale bar = 10µm. For A-C, N = 7 animals, n ≥ 25 nuclei/treatment. (D) Gene expression analysis showed significant increased p53 and p21 in stretch injury CFs. N = 9 animals. Error bar = sem. Linear mixed model, ANOVA, **p<0.01, *p<0.05.

We further investigated gene expression for markers of cell cycle arrest, p16, p21, p53, in the stretched cultures via RT-qPCR studies. The p53-p21-RB axis is known to be involved in DNA damage response signaling and cell cycle arrest (*43*, *44*). Stretch Injury CFs had increased expression of p53 and p21 but surprisingly had decreased expression of p16 (**Fig. 2D**). Together, we found mechanical perturbation of primary murine cardiac CFs induced a senescent-like phenotype when assessed through markers of cell cycle, nuclear morphology, and DNA damage.

### Chemical and mechanical induction of senescence have unique pathways

Given that a combination of markers is necessary to define a senescent phenotype and several markers are not senescent exclusive, we sought to additionally induced senescence through oxidative stress, reactive oxygen species (ROS)-induced senescence, which is observed after cardiac injury and in cardiovascular disease, especially after injury (*40*). We cultured primary WT CFs over 21 days with hydrogen peroxide treatment to chemically induced senescence (**Fig. 3A**). We then assessed ROS-induced CFs for the same markers as the mechanically-induced CFs. We found the ROS-induced CFs had a significant decrease in Ki-67 staining compared to TCP controls indicating decreased proliferation. Using the same quantification of lamin B ring thickness, we found a decrease in ring thickness (**Fig. 4A**) in ROS-induced CFs and general decrease in lamin B intensity as shown in the representative images. Analyzing morphological changes, we observed a significant increase in nuclear area in ROS-induced CFs (**Fig. 4B**). Interestingly, while we observed decreases in the aspect ratio and chromatin condensation, this is opposite to the phenotype detected with the mechanically-induced senescent CFs where we observed increased aspect ratio and chromatin condensation (**fig. S2B** and **C**).

**Figure 3.**
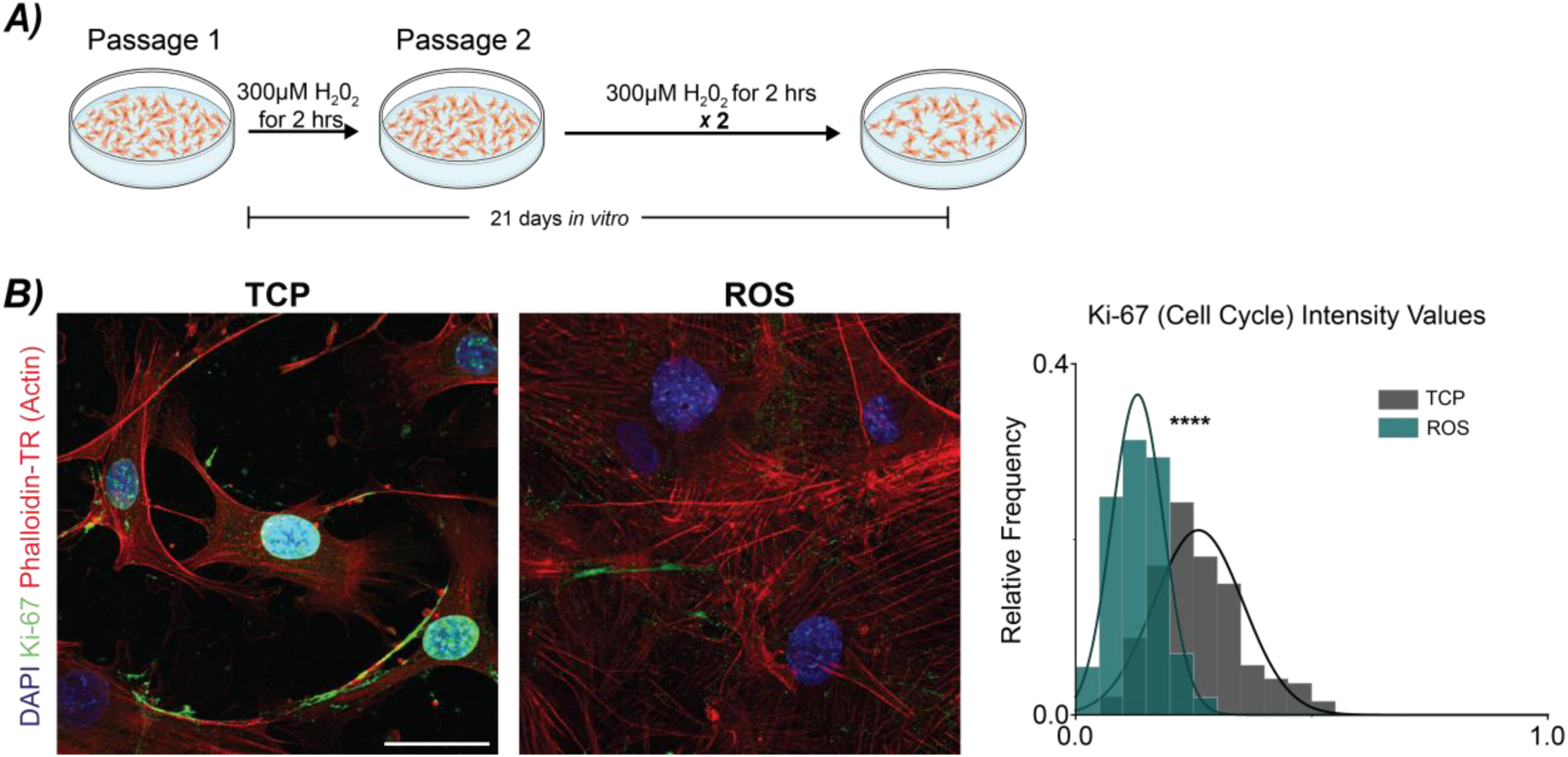
Validating markers of mechanically-induced senescence through reactive oxygen species (ROS)-induced senescence. (A) Isolated CFs from 4-9 week old mice were plated on TCP and treated with 300µm H_2_0_2_ for 2 hours over the course of 21 days *in vitro* to induce a senescent phenotype. (B) Representative images show Ki-67 staining of CFs in TCP and ROS cultures. Quantification of Ki-67 positive cells in TCP and ROS cultures indicated a decreased relative frequency of cycling cells in ROS cultures, noting a decrease in proliferation. DAPI = blue (405nm), Ki-67 = green (488nm), Actin (Phalloidin-TR) = red (561nm). N = 7-8 animals, n ≥ 25 nuclei/animal/treatment. Scale bar = 50µm. Linear mixed model, ANOVA. ****p <0.0001.

**Figure 4:**
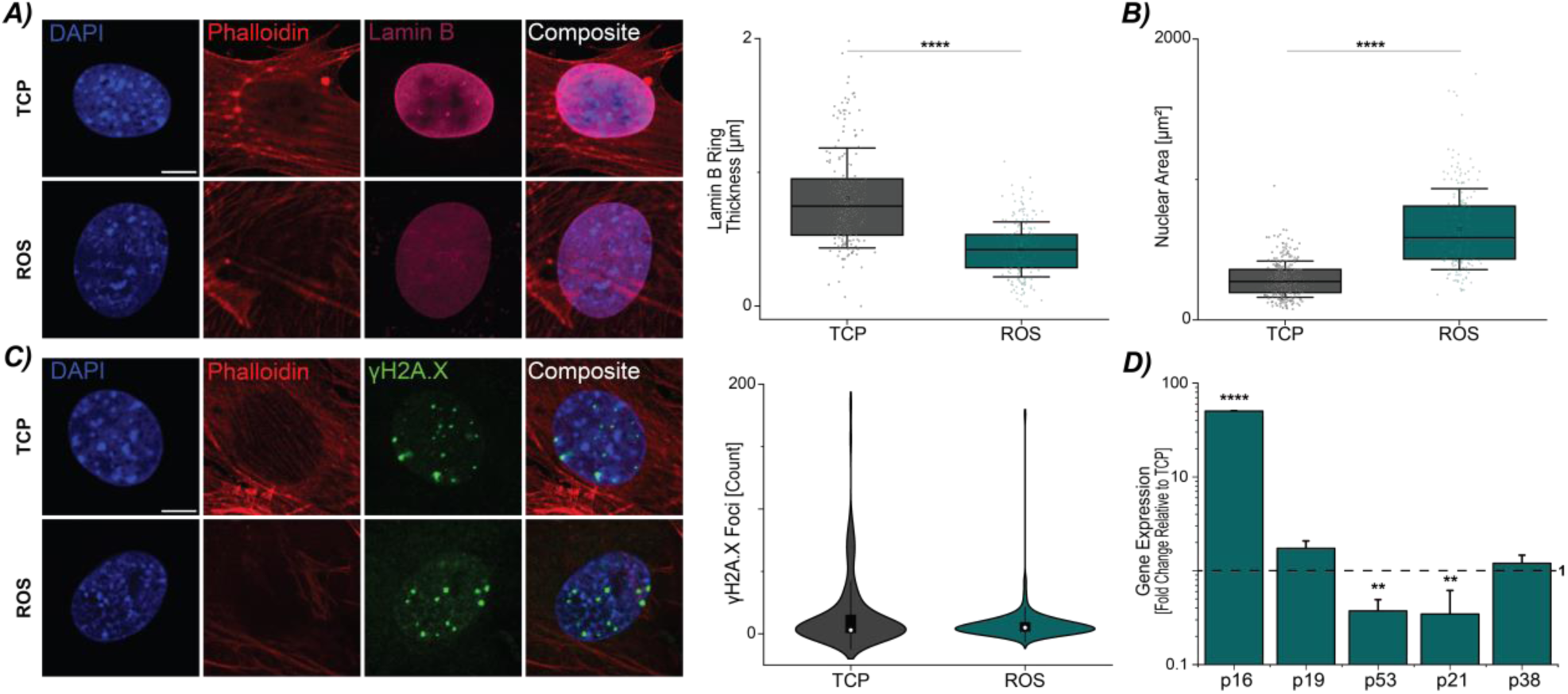
Cardiac fibroblasts experiencing oxidative stress (ROS) display markers indicative of a chemically-induced senescent phenotype, including alterations in morphology, DNA damage, and gene expression. (A) Representative images of lamin B staining from CFs in TCP and ROS cultures. Cells in ROS cultures showed a decrease in the lamin B ring thickness. Scale bar = 10µm. Error bar = 1 Std. N = 7-8 animals, n ≥ 25 nuclei/treatment. (B) ROS-induced CFs had an increase in nuclear area. Error bar = 1 Std. N = 8-10 animals, n ≥ 25 nuclei/treatment. (C) Representative images from TCP and ROS cultures showing ɣH2A.X foci. Nuclei from CFs in ROS cultures did not show increased ɣH2A.X foci compared to TCP CFs. Error bar = 1 Std. Scale bar = 10µm. N = 6-8 animals, n ≥ 25 nuclei/treatment. (D) Gene expression of ROS cultures has increased p16 expression, and decreased p53 and p21 expression. N = 8 animals. Linear mixed model, ANOVA. ****p<0.0001, **p<0.01.

We examined the ROS-induced senescent CFs for increased DNA damage through ɣH2A.X foci count, and we found no difference between TCP and ROS-induced CFs (**Fig. 4C**). The lack of a significant difference in ɣH2A.X foci was supported by gene expression analysis for p53 and p21 in which a decrease in both genes were observed compared to TCP controls (**Fig. 4D**). However, ROS-induced CFs showed increased p16 expression. Taken together, the mechanically-induced senescent-like CFs display similar hallmarks of senescence as ROS-induced CFs with decreased proliferation (i.e., Ki-67), decreases in lamin B, and increases in nuclear area. However, we found the modes of induction could be unique and dependent on the biomechanical vs biochemical stimulus. While both showed increased gene expression in markers of cell cycle arrest, ROS-induced CFs showed significant increases p16 expression and mechanically-induced CFs had increases in p53 and p21 expression.

### Loss of nuclear structural integrity does not enhance the initiation of a senescent state

Given that many alterations and markers found in senescent cells also overlap with modifications in the nucleus and nuclear envelope, we hypothesized that mechanically-sensitive proteins in the nuclear envelope might play a role in the initiation of the senescent-like phenotype observed in the WT stretch cultures. We used a mouse model where the lamin A/C protein in the nuclear lamina is modified so that the lamin A/C protein does not integrate with the nuclear envelope (*45*). Lamin A/C maintains the mechanical tension of the nuclear envelope and has been shown to interact with p16 and p53 (*46*). We again established primary cultures of CFs from mice between the ages of 4-9 weeks as mice with the mutation (partial Lamin A/C knockout) are severely runted and die of cardiac failure around 9 weeks. Next, we evaluated the cultures under the same two loading conditions, Stretch Control and Stretch Injury. We first assessed the cultures for Ki-67 expression. We observed a significant decrease in the Ki-67 distribution in Stretch Injury CFs compared to Stretch Control CFs (**Fig. 5A** and **fig. S3A**), and similar to what we observed in **Fig.1A**. Examining the Stretch Control and Injury partial lamin A/C knockout (lamin A/C null) CFs for the defined morphological markers of senescence, we observed no change in nuclear area (**Fig. 5B** and **fig. S3B**) or aspect ratio (**fig. S3E**). However, we found an increase in chromatin condensation (**Fig. S3D**), similar to the WT mice (**Fig. S1B**). Interestingly, Stretch Injury lamin A/C null CFs had increased lamin B ring thickness compared to Stretch Controls (**Fig. 5C**), possibly as a compensatory mechanism in absence of lamin A/C. While we did not observe a decrease in lamin B ring thickness which would confirm a senescent-like phenotype, we noted an increase in nuclear rupture and regions of lamin B dilution in the mechanically-stimulated (Stretch Control and Injury) lamin A/C null CFs suggesting decreased structural integrity by the loss of lamin A/C along the nuclear envelope. This rupture occurred at the site of lamin B dilution but not at the highest area of curvature within the CF nucleus (**fig. S4A**) (*47*). On examining the DNA damage response, we observed increase ɣH2A.X foci in the Stretch Injury lamin A/C null CFs (**Fig. 5D**). However, gene expression analysis did not show significant difference in p53 or p21, which were both upregulated for the WT mechanically-induced senescent CFs (**fig. S3B**).

**Figure 5:**
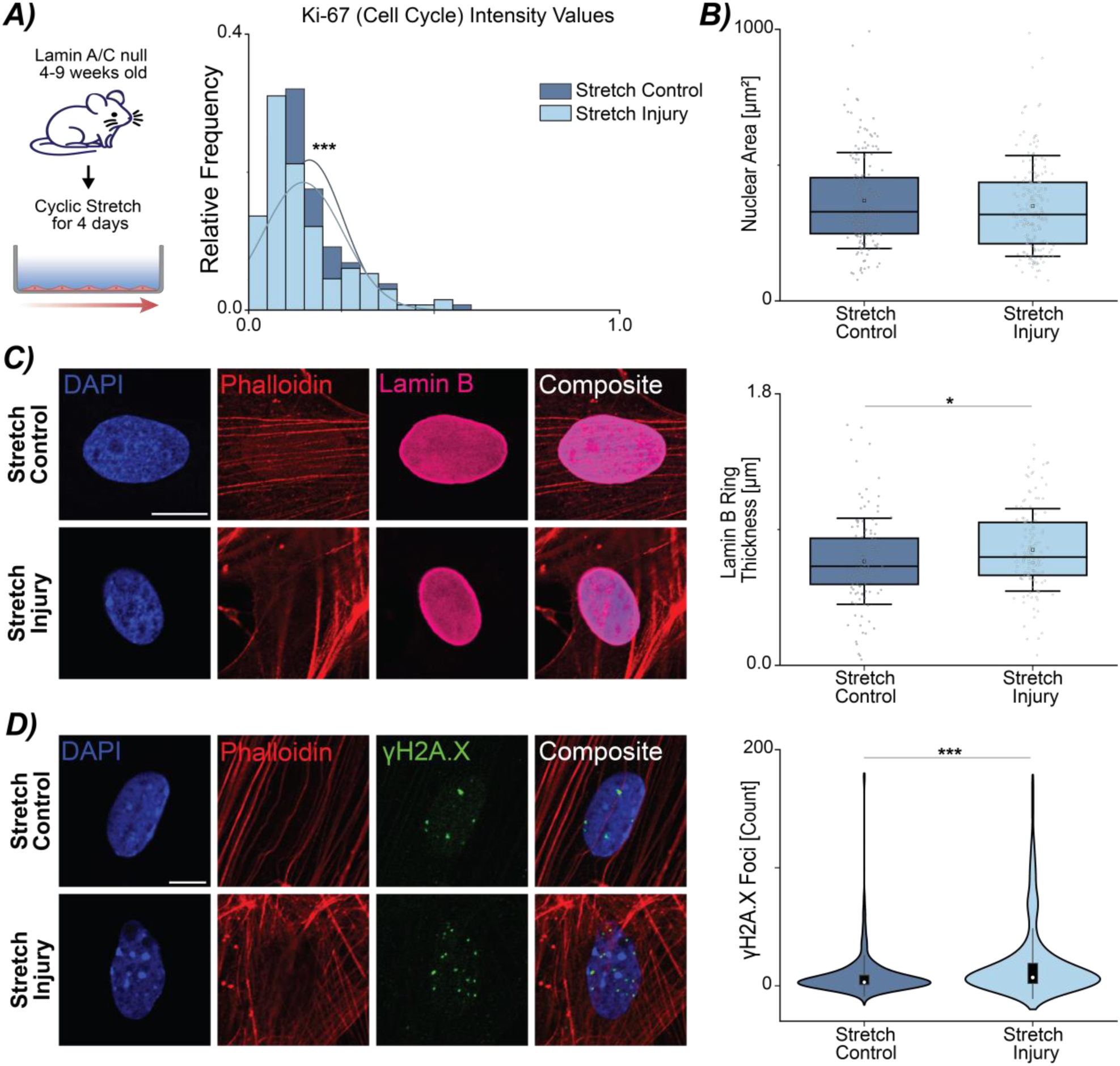
Disruption of nuclear integrity through loss of Lamin A/C does not amplify a premature, mechanically-induced senescent phenotype in cardiac fibroblasts. (A) Isolated CFs from 4-9 weeks old lamin A/C null mice subjected to stretch injury have decreased Ki-67 positive cells compared to stretch control cells. N = 5 animals. (B) Mechanical stretch does not alter nuclear area in Stretch Injury CFs compared to Stretch Control CFs. N = 7 animals. (C) In lamin A/C null CF nuclei, Stretch Injury loading results in an increase in lamin B thickness compared to stretch control. N = 5 animals. (D) Stretch injury CFs have an increase in ɣH2A.X foci. N = 7 animals. Scale bar = 10µm. B-D: Error bar = 1 Std. A-D: n ≥ 25 nuclei/treatment. Linear mixed model, ANOVA. ***p<0.00, *p<0.05.

Recognizing that due to the interaction between lamin A/C and p16 (*46*), a senescent phenotype may again be unique, we performed the hydrogen peroxide treatment of the lamin A/C null CFs to generate a ROS-induced senescent phenotype. Besides a decrease in lamin B thickness, ROS-induced lamin A/C null CFs displayed similar hallmarks of senescence as WT CFs (**fig. S5**). This demonstrated the necessity for describing several markers to characterize the senescent phenotype, as lamin B ring thickness did not decrease in the mechanically- or chemically-induced lamin A/C null CFs. Take together, the compilation of morphological readouts, gene expression, and DNA damage did not indicate the progression of a premature senescent state with altered nuclear structural integrity through the loss of lamin A/C.

### Mechanically-sensitive protein, Emerin, decreases with Stretch Injury

Since we did not observe an exaggerated phenotype with the lamin A/C null CFs, we looked at other mechanically-sensitive proteins in the nuclear envelope. Emerin is a nuclear envelope protein which tightly interacts with lamin A/C in regulation nuclear mechanics (*48*, *49*). Studies have found that increased strain can cause emerin to locate to the outer nuclear membrane(*50*). Using the same stretching protocol to initiate a mechanically-induced senescent phenotype (**Fig. 1B**), we next examined emerin by immunofluoresence. We found Stretch Injury CFs had a significantly decreased emerin ring thickness compared to Stretch Control CFs (**Fig. 6A**). Curious if this finding was maintained by all senescent CFs, we quantified the emerin ring thickness in ROS-induced and TCPs CFs and did not find alterations with chemically-induced senescent CFs (**Fig. 6B**). In all lamin A/C null treatments, mechanically- and chemically-stimulated, we measured decreased emerin ring thickness but no significant differences between the different treatments (**fig. S6A and B**). This is not unexpected as emerin localization in the lamin A/C null CFs has been shown to be largely localized to cytoplasm (*45*). Additionally, with the decreased emerin ring thickness, we also qualitatively observed irregular actin structural patterns immediately surrounding the nuclear envelope. The significant difference in WT mechanical-induced senescent CFs compared to Stretch Controls demonstrated the importance of nuclear structural integrity and its interplay with the induction of a senescent phenotype.

**Figure 6:**
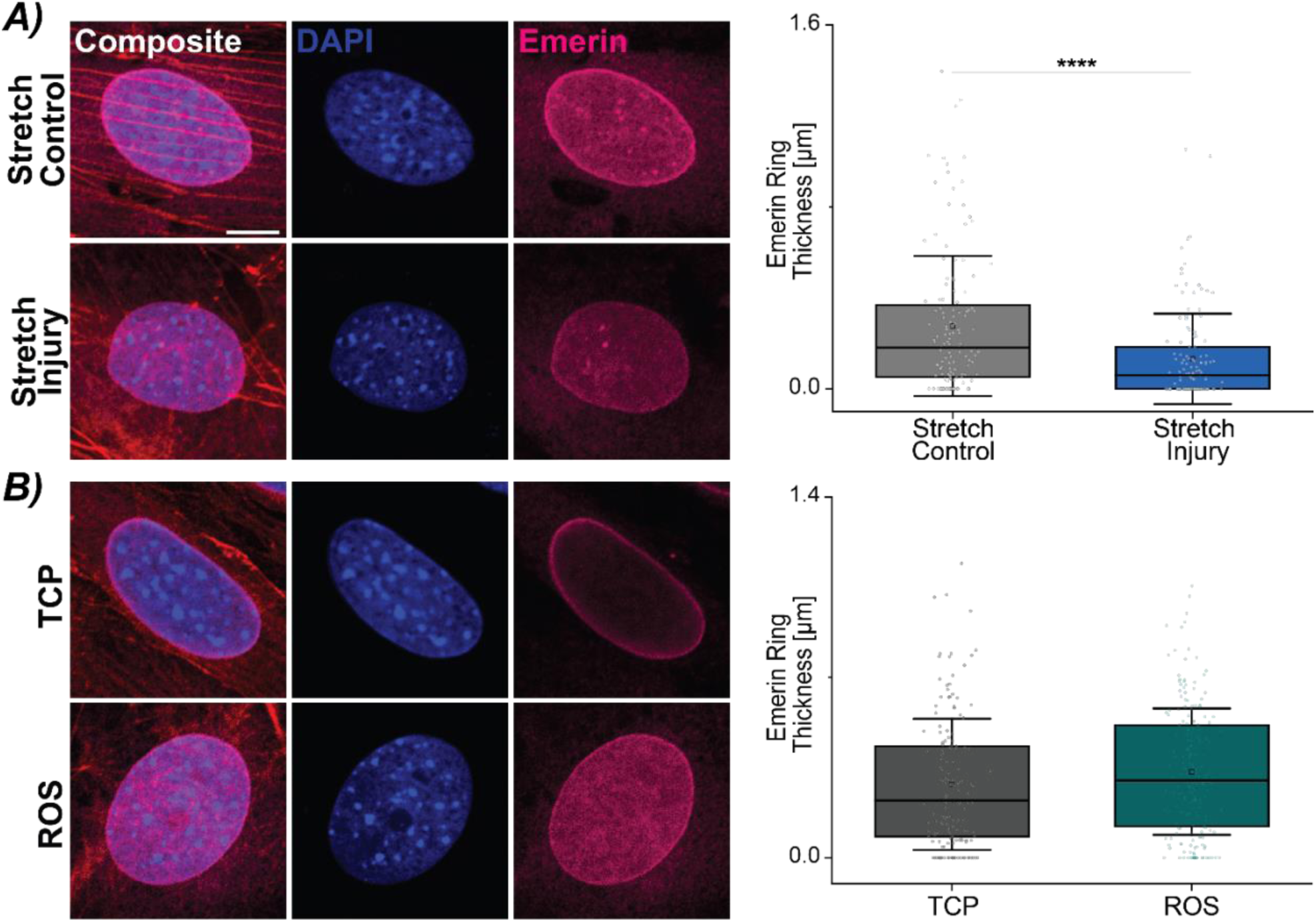
Mechanosensitive nuclear envelope protein, emerin, may be a key mediator in the initiation of a mechanically-induced senescence phenotype in cardiac fibroblasts. Representative images show emerin staining for nuclei from WT CFs from mechanically-induced, chemically-induced, and respective control cultures. Scale bar = 10µm. (A) Quantifying the emerin ring thickness in Stretch Control and Stretch Injury CFs indicated a decreased ring in Stretch Injury CFs. N = 5 animals. (B) Emerin ring thickness between TCP and ROS-induced CFs showed no significant alterations with the induction of a senescent phenotype. N = 6-8 animals, n ≥ 25 nuclei/treatment. Error bar = 1 Std. Linear mixed model, ANOVA. ****p<0.0001.

## Discussion

In this study, we examined how perturbed mechanical stimulus can aid in the initiation of cellular senescence in a simplified *in vitro* model of cardiac disease. Using previously established markers of cellular senescence, we found that perturbed mechanics can alter proliferation, lamin B expression, induce a DNA damage response, and increases in p21 and p53 expression to define a mechanically-induced senescent profile in CFs. Furthermore, we found mechanically-sensitive proteins in the nuclear envelope, emerin, maybe involved at the initiation state.

### Mechanical Stimulation and Cellular Senescence

With continued growth in the fields of aging and cellular senescence, the factors which induce the transition towards senescence phenotype are becoming more well defined. This adds to the complexity in defining this cellular state as there is still no single marker of cellular senescence (*10*). Instead, numerous markers which encompass both morphological and biochemical responses are necessary to determine the phenotype (*6*). Additionally, the markers associated with the induction of cellular senescence can vary within the same cell type, depending on the type of stimulation and the activated pathway. This heterogeneity occurs even at a single cell level (*6*, *44*, *51*). Among the various mechanisms being explored, mechanical stimulation is increasingly being recognized as a key factor in triggering cellular senescence.

In diseases such as intervertebral disc degeneration and osteoarthritis, increases in dynamic cyclic loading using nucleus pulposus cells or chondrocytes have been shown to induce premature senescence through DNA damage response signaling and the p53/p21 axis, not p16 (*11*, *52*), similar to our results. Interestingly, in the same cell type, nucleus pulposus cells, high magnitude compression also initiated cellular senescence indicating abnormal mechanical loading can initiate this phenotype (*53*). Additionally, studies have examined how the stiffening of the local microenvironment can trigger alterations (*12*). Taken together, these studies demonstrate that perturbed mechanical stimulation can be one of the initial factors in the transition. However, in these studies, a significant increase in strain magnitude (at least 4-fold higher) or overloading of the cells is frequently used to stimulate a senescent state (*11*, *12*). In contrast, our results showed a decrease in (hypophysiologic) strain magnitude and increased cyclic frequency as a key driver for the senescent transition indicating that maybe any alteration in cellular mechanics outside of ‘normal’ physiological loading could initiate senescence.

We were surprised to observe that senescence was initiated through the activation of distinct pathways, resulting in differences in the cellular senescent phenotype. Comparing WT Stretch Injury CFs to those activated through oxidative stress (ROS-induced) showed two unique profiles indicating differing pathways of activation. ROS-induced CFs had increased expression in p16, but we observed no differences in DNA damage through ɣH2A.X compared to CFs on TCP. Stretch Injury CFs had increased p21 and p53 expression, and increased DNA damage response compared to Stretch Control CFs. Both modes of induction had decreases in lamin B ring thickness and Ki-67 expression. Similarly, lamin A/C null mechanically-stimulated CFs and ROS-induced CFs showed differing phenotypes. Again, these cultures differed in gene expression and lamin B expression. In addition, the cultures also differed in lamin B expression measured using microscopy images and quantification of lamin B ring thickness. This difference could be explained by the mechanical stimulation the Stretch Injury CFs experience. The increase in lamin B may be a compensatory mechanism for the lack of lamin A/C which is known to regulate nuclear tension (*54*, *55*).

### Nuclear Integrity and Mutations in Lamin A/C

Lamin A/C is a nuclear envelope protein that supports the nucleus both structurally in maintaining both shape and size, but additionally interacts with genes necessary for cell cycle and DNA damage repair. There is an array of diseases associated with the mutations in the *Lmna* gene, termed laminopathies. Depending on the mutation or deficiency, phenotypic effects range from diseases which cause accelerated aging as seen in Hutchinson-Gilford Progeria Syndrome, to those which result in severe retardation of skeletal muscle and cardiac disease as observed in Emery-Dreifuss muscular dystrophy (EMDM) (*39*). The large-scale effects resulting from disruption in the nuclear envelope show the importance of the nuclear envelope and its integrity for healthy cell and tissue maintenance (*56*). A complete deficiency in lamin A/C in the nuclear periphery has been shown to lead to increased nuclear fragility, altered chromatin organization, nuclear rupture, blebbing and increases in DNA damage (*54*, *57*, *58*). As lamin A/C helps to regulate the nuclear tension, it has also been shown to be mechanically sensitive to both gradual changes in the microenvironment and applied forces (*8*, *54*). In this study, we found that alterations in the nuclear envelope, specifically with the knockout of lamin A/C, led to increased DNA damage within the nucleus and severe nuclear ruptures within the stretched cultures. However, in the mechanically active CF cultures, nuclear blebbing was not observed, which was observed in the CFs plated on TCP and ROS-induced CFs, and has been described in the literature (*55*). Interestingly, an increase in lamin B thickness was observed in lamin A/C null CFs including CFs with nuclear rupture. In the lamin A/C null cells, lamin B may additionally act as a compensatory mechanism in nuclear maintenance of both shape and size.

Lamin A/C has been shown to interact with p16 and disruption of this interaction alters the cell cycle (*46*). In mechanically stretched cultures, lamin A/C null CFs did not show increased p53/p21 or p16 even though a decrease in Ki-67 was observed. It is unclear if gene expression for cell cycle arrest occurred or if there was increased cell death through necrotic or apoptotic pathways, leading to the decreased proliferation observed. Future experiments looking at apoptosis in the lamin A/C null CFs and increased mechanical-stimulation could be performed to clarify the difference observed between gene expression and Ki-67 staining.

### Nuclear Integrity and Emerin

Within the nuclear envelope, other proteins have been identified as mechanically sensitive, for example, the linker of nucleoskeleton and cytoskeleton (LINC) complex and emerin (*8*, *49*, *59*). Disruption in emerin also has large scale effects in development and is another known cause of muscular dystrophy. Primarily located on the inner nuclear membrane, emerin transfers information between the cytoskeleton to the nucleus. Moreover, emerin interacts with not only with the nuclear envelope proteins, but also interacts with lamin A/C and lamin B, and binds to actin to maintain chromatin architecture, nuclear shape and aspect ratio (*48*, *49*, *60*). Disruption or complete knockdown of emerin shows changes in F-actin patterning and impaired actin binding (*48*, *60*). Alterations in strain and actin polymerization have been noted following stretch injury (*61*), and separately have shown increases of emerin in the outer nuclear membrane (*50*). Our results agree with these findings showing alterations in the localization of emerin within the nuclear envelope and cytoplasm of the CF. The lack of an inner emerin ring in the lamin A/C null cultures, both mechanical and chemical, and Stretch Injury CFs indicates that emerin acts as a key element in maintenance of strain transfer with active mechanical forces. Future studies could examine emerin’s interaction with actin and localization within senescent CFs as potential therapeutic target. Indeed, regulation of the nuclear envelope has already been implicated in mechanically-stressed induced senescence in skeletal muscle MSCs which indicates a significant connection between induction of cellular senescence and nuclear integrity (*62*).

From our simplified *in vitro* model of the mechanical perturbations observed in cardiac disease, we found altered mechanical stimulation in CFs leads to a mechanically-induced premature senescent-like phenotype. The study demonstrates how hypophysiologic changes in mechanical stimulation, and more specifically, a decrease in strain and increased frequency can initiate cellular senescence. Additionally, we identify a set of markers to characterize a mechanically-induced phenotype advancing the effort to more precisely define cellular senescence.

## Materials and Methods

### Cardiac Fibroblast Cultures

Primary murine cardiac fibroblast (CF) cultures were derived from B6.129S1(Cg)-*Lmna^tm1Stw^*/BkknJ wild type (WT) and null (Mut) mice (Jackson Labs, Stock No.: 009125) (*45*). The study included mice between the ages of 4-9 weeks with no exclusion of sex. Specific pathogen-free, temperature-controlled housing with 12-hour light cycles maintained the mice, which received food and water *ad libitum*. All animal procedures followed University of Colorado Boulder Institutional Animal Care and Use Committee approval. Cardiac tissue was excised from mice and placed immediately in cold DPBS (Hyclone, SH30028.03). A 25-gauge needle inserted in the apex of the tissue allowed flushing of hearts with 30mL of cold DPBS. The tissue was then minced with a scalpel and washed three times in ice cold HBSS (Gibco, 4500-462) in a sterile environment. Digestion medium, containing 0.2% collagenase P (Roche, 11213873001) in DMEM supplemented with 3% FBS (Gibco, 261-40-079) and 10 µg/ml DNase1 (Sigma, D4263-5VL), was added to the minced tissue. The tissue underwent digestion for 25 mins at 37°C at 900 rpm. Following this period, the tissue was triturated ten times with a P1000 tip to mechanically disrupt the tissue. The digested mixture was incubated for an additional 10 mins at 37°C at 900 rpm. After a second mechanical disruption, the tissue was filtered through a 40µm cell strainer and rinsed with supplemented medium, D10. The cardiac fibroblast medium, D10, consisted of DMEM/F12 (Gibco, 1330-032) supplemented with 10% FBS (Gibco, 261-40-079), 1× Penicillin/Streptomycin (Gibco, 15140-122), and 1µM ascorbate-2-phosphate (Sigma, 49752-10G). Addition of 1mM EDTA (Sigma, 03690-100) to D10 quenched the digestion reaction. The resulting cell/tissue pellet was lysed with 1× RBC Lysis Buffer (ebioscience, 00-4300-54) and the reaction quenched with D10. We plated the cells in 100 mm dishes for 2.5 hours in a 37°C incubator with 5% CO_2_. After 2.5 hours, we washed petri dishes three times in warmed PBS and added D10 back to the dishes. Based on previously established methods of isolation and characterization, we deemed the remaining cells adherent to the petri dish cardiac fibroblasts (*63*).

Passage 1 CFs were divided up into 4 uses: RNA, mechanical stimulation, immunofluorescence (e.g., TCP, ROS, mechanical), and continued culture. For tissue culture plastic (TCP) controls for RNA and immunofluorescence (IF), cells were plated at a density of 5 × 10^3^/cm^2^ in 8 well ibidi (IF) dishes and 6 well plates (RNA). For mechanical stimulation experiments, both passage 1 and passage 2 cells were used. All CFs used in mechanical stimulation experiments were plated at 6 ×10^3^ cells/cm^2^ on a MCFX 16 well flexible membrane. Prior to plating, we coated the membranes with Geltrex (Gibco, A1569601) for 16 hours at 37°C. For continued culture CFs, passage 1 CFs were plated at 2.5 × 10^3^ in 100mm petri dishes and cultured for 5 days with medium changes every 2 days. On day 5, cells were detached using TrypLE, quenched with D10, and pelleted at 1300 rpm. CF pellets were resuspended in D10 and filtered through 70µm cell pellet prior to count with Trypan Blue. We divided the CFs into groups for experimental use IF, RNA, and mechanical stimulation. The same coating protocol of Geltrex and plating density was used for P2 CFs subjected to mechanical stimulation. After 4 days of mechanical stimulation, cells were used for RNA and IF. For IF experiments, cells on MCFX plates were fixed in 4% PFA in DPBS for 20 mins at room temperature. After 20 mins, fixed cells were washed 3 times in DPBS and stored at 4°C in DPBS until use.

For TCP controls, cells were plated at 5 × 10^3^/cm^2^ in a 6 well plate. All TCP controls for imaging and IF were collected 3 days after plating. TCP cells used in IF experiments were fixed in 4% PFA in DPBS at room temperature for 10 mins. After 10 mins, fixed cells were washed in DPBS and stored in at 4°C in DPBS. For all experiments, RNA was collected by lysing the CFs in Qiazol (Qiagen, 79306) and stored at −80°C until extraction.

### Mechanical Stimulation Experiments

We used Passage 1 and Passage 2 CFs for all mechanical stimulation experiments. After plating CFs in the flexible 16 well-membranes, cells were allowed to adhere for 24 hours at 37°C in 5% CO_2_ before stretching.

### Mechanical Stimulation

Two loading modes were designed: (1) Stretch Control and (2) Stretch Injury. Using a MechanoCulture FX (CellScale, MCFX) device, we subjected cells to uniaxial stretch at different strain magnitudes and rates. The Stretch Control loading scheme was continuously stretched for 4 days at 8.5% strain at a frequency of 1 Hz within an incubator maintained at 37°C in 5% CO_2_. The Stretch Injury loading scheme was 4 days total with 24 hours of mechanical stimulation at 8.5% strain at 1 Hz and 72 hours at 2.5% strain at 2 Hz within an incubator maintained at 37°C in 5% CO_2_ (Fig 1A). Full medium changes were performed every 24 hours to decrease any confounding biochemical effects.

### ROS Treatment

Passage 1 cells were plated at 4 × 10^3^/cm^2^ in 3 wells of a 6 well TCP dish. After 24 hours, cells were treated with 300µM H_2_0_2_ (J.T. Baker, 2186-01) for 2 hours at 37°C (*64*). After 2 hours, wells were washed one time with D10 and cultured in D10 for 48 hours. Minimal cells death was observed after the first treatment. After 48 hours, we passed the CFs, and all remaining treatments were performed without further passaging. Cells were divided up between RNA and IF experiments after the first treatment. For RNA collection, CFs were plated at 4 × 10^3^/cm^2^ in a 6 well TCP plate. For IF, CFs were plated at 4 × 10^3^/cm^2^ in 3.5cm^2^ imaging dishes (ibidi, 50-305-806). WT ROS-induced CFs received a total of three treatment of 300µM H_2_0_2_ for 2 hours at 37°C. Lamin A/C null ROS-induced CFs received a total of two treatment of 300µM H_2_0_2_ for 2 hours at 37°C due to increased cell death observed in the cultures. Culture medium was changed every 3-4 days. RNA and IF dishes were collected 8 days after the final H_2_0_2_. During the 8-day incubation period, increased cellular and nuclear area was observed indicating potential initiation of cellular senescence. These CFs are referred to as ROS-induced CFs in the results section.

### RNA Isolation and RT-qPCR

RNA was isolated using Direct-zol RNA Miniprep (Zymo, 76020-642), nanodropped for concentration and quality, and reverse transcribed into cDNA using iScript^TM^ Reverse Transcription Supermix (BioRad,1708841). Primers were designed using NCBI primer BLAST with all primers designed to span an exon-exon junction (Table S1). Real-time quantitative PCR was performed with SsoAdvanced^TM^ Universal SYBR^®^ Green Supermix (BioRad, 1725271) in a CFX96 Touch™ thermocycler (BioRad) using 10 ng of cDNA/reaction. All data was normalized to *Gapdh* prior to fold change. The relative change in gene expression was quantified using the ΔΔCt method.

### Immunofluorescence and Imaging

For immunofluorescence, fixed CFs were permeabilized in 1% Triton-X in PBS and blocked with 1% normal goat serum with 1% BSA in PBT (0.125% Tween 20 in PBS) prior to staining. CFs were stained with primary antibodies in antibody buffer (1% BSA in PBT) overnight at 4°C rocking and washed 3 times in PBT to remove any unbound antibody. CFs were incubated in secondary antibodies for 45 mins at room temperature. Following two washes in PBT, CFs were stained for DAPI (1:1000; Invitrogen, D1306) and Phalloidin Texas-Red (1:300, Life Technologies, T7471) for 30 mins. ROS CFs and TCP CFs were stored at 4°C in PBS until imaged. Due to the thickness of the flexible membrane of the MCFX device, wells were punched out of the flexible plate, flipped, and mounted in DAKO Mounting Medium (Agilent, S302380-2) for improved resolution for imaging. Primary Antibodies: Mouse anti-Mouse ɣH2A.X (1:400, CST, 80312S), Rabbit anti-Mouse Lamin B1 (1:500, abcam, ab16048), Rat anti-Mouse Ki-67 (1:300, ebiosciences, 14-5698-82), Rabbit anti-Mouse Emerin (1:250, CST, 30853S). Secondary antibodies: Goat anti-Mouse IgG Alexa Fluor 488 (1:500, ThermoScientific, A28175), Goat anti-Rabbit IgG Alexa Fluor 633 (1:500, ThermoScientific, A21070), Goat anti-Rat IgG Alexa Fluor 488 (1:100, BioLegend, 405418).

For imaging acquisition, all fixed and mounted wells were imaged on a Nikon A1R Confocal using 512 x 512 pixel field of view with a 60X oil immersion objective. Greater than 25 nuclei per treatment and genotype were taken. For imaging analysis, a custom MATLAB code analyzed the collected images of the cell nucleus. The process involved cropping, segmenting, and histogram normalizing images for biophysical features. Calculations of lamin B and emerin ring thickness relied on detecting an outer ring around the nucleus based on fluorescence intensity and subtracting it from the nuclear area. MATLAB’s built-in functions determined other biophysical features such as nuclear area. Detection of ɣH2A.X foci employed an intensity threshold method, counting peaks above the set threshold. Proliferation analysis with Ki-67 involved histogram normalized intensity values examined against relative frequency.

### Statistical Analysis

We analyzed the difference between treatments (TCP and ROS or Stretch Control and Stretch Injury) for lamin B Ring, emerin ring, Ki-67, gene expression analysis, and ɣH2A.X foci count using linear mixed effects models (nlme package, Version 3.1-140) in R (RStudio, Version 1.2.1335; R, Version 3.6.1, Boston, MA). Animal was considered a random effect in the models. Type II Sum of Squares ANOVA tested differences between treatments. For ɣH2A.X foci data, we analyzed the model with and without the outliers to confirm statistical significance and p-values. Post-hoc testing utilized the emmeans package with Tukey’s HSD corrections for multiple comparisons (Version 1.4.3.01). ANOVA normality assumptions underwent validation through testing model residuals with the Shapiro-Wilk test and visual examination of qq-plots. If necessary, data was transformed to meet the normality assumptions of ANOVA.

## Acknowledgments

SES is grateful for support from the NIH T32 GM-065103.

## Author contributions

Conceptualization: SES, CPN.

Methodology: SES, AKS, EYM, SG.

Investigation: SES, AKS, KG.

Visualization: SES, EYM, SG.

Supervision: SES, CPN.

Writing—original draft: SES, CPN

Writing—review & editing: SES, AKS, KG, EYM, SG, CPN.

## Data and materials availability

All data needed to evaluate the conclusions in the paper are presented in the paper or Supplementary Materials. Code available upon request.

---

**Fig. S1.**
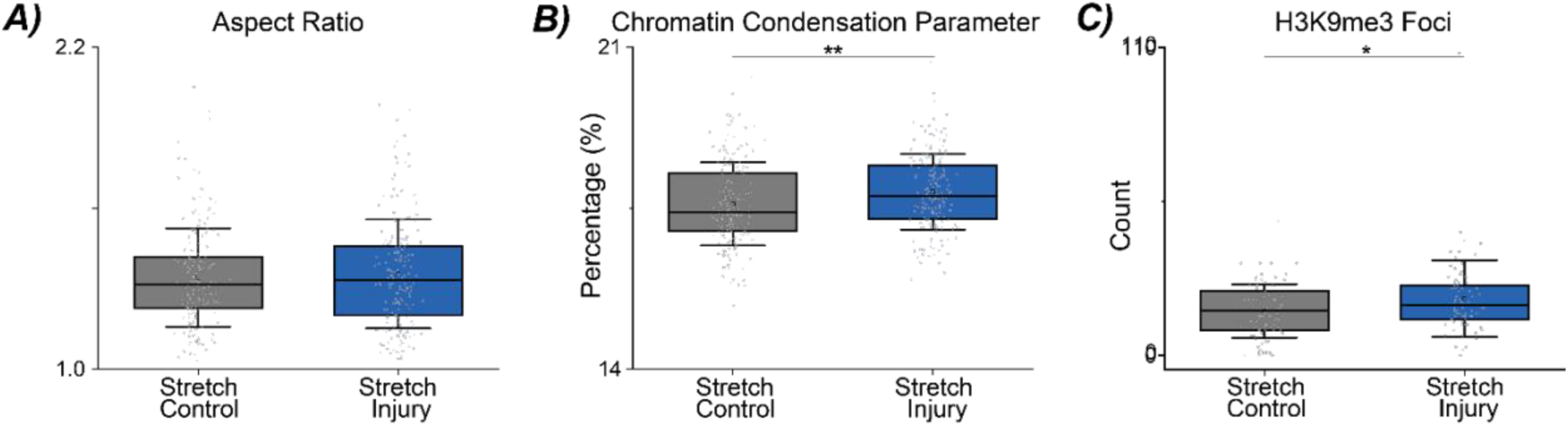
Analysis of the morphological features for mechanically-induced senescent cardiac fibroblasts (CFs) (A) A difference in aspect ratio was not observed when comparing stretch control and stretch injured CFs. N = 8-10 animals, n ≥25 nuclei/treatment. (B) Alterations in DNA condensation were observed and quantified using Sobel edge detection in MATLAB. Stretch injured CFs had significantly increased chromatin condensation compared to stretch controls. N = 8-10 animals, n ≥ 25 nuclei/treatment. (C) With increased chromatin condensation, CFs were stained for H3K9me3 in both Stretch Control and Stretch Injury treatment groups. Stretch Injury CFs had a slight increase in H3K9me3 foci compared to Stretch Controls. N = 3 animals, n ≥ 25 nuclei/treatment. Error bar = 1 Std. Linear mixed model, ANOVA. **p<0.01, *p<0.05.

**Fig. S2.**
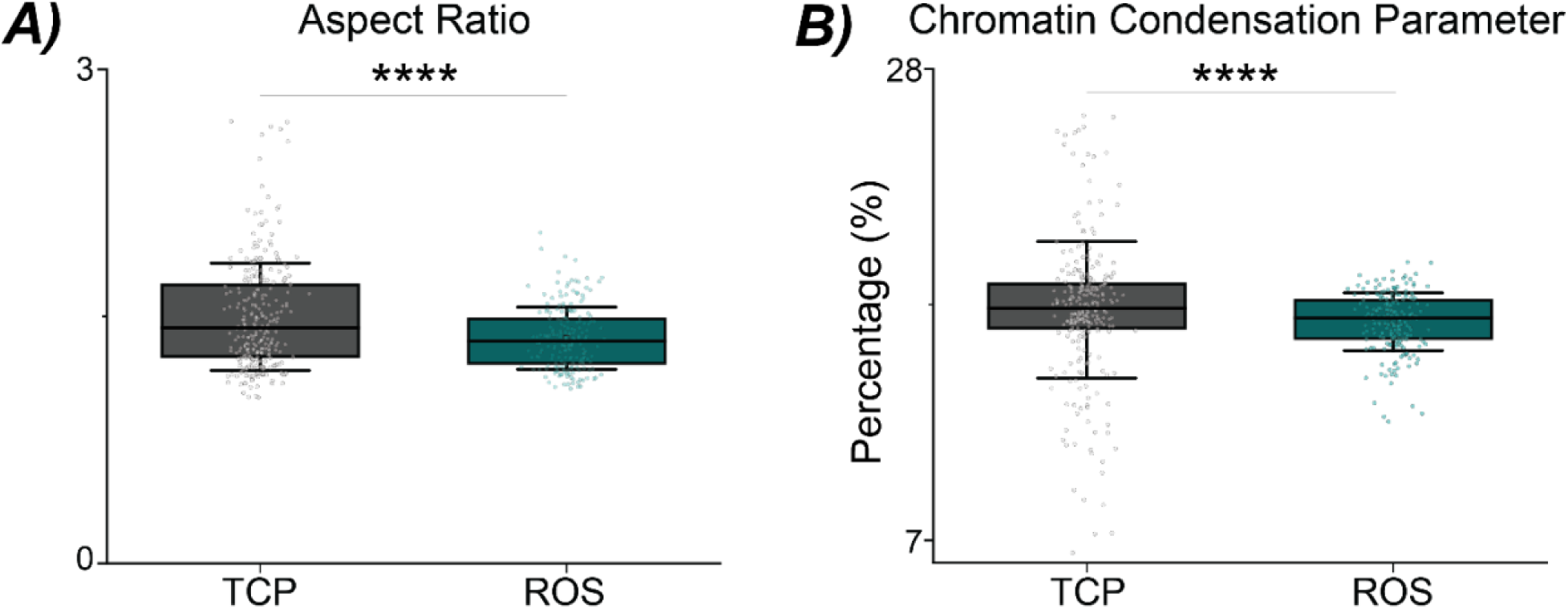
ROS-induced senescent CFs show alterations in nuclear shape and chromatin condensation. (A) Senescent CFs show a significant difference in aspect ratio compared to CFs plated on TCP. (B) Compared to the control TCP CFs, ROS-induced senescent CFs have decreased chromatin condensation. N = 8-10 animals, n ≥ 25 nuclei/treatment. Error bars = 1 std, Linear mixed model, ANOVA. ****p<0.0001.

**Fig. S3.**
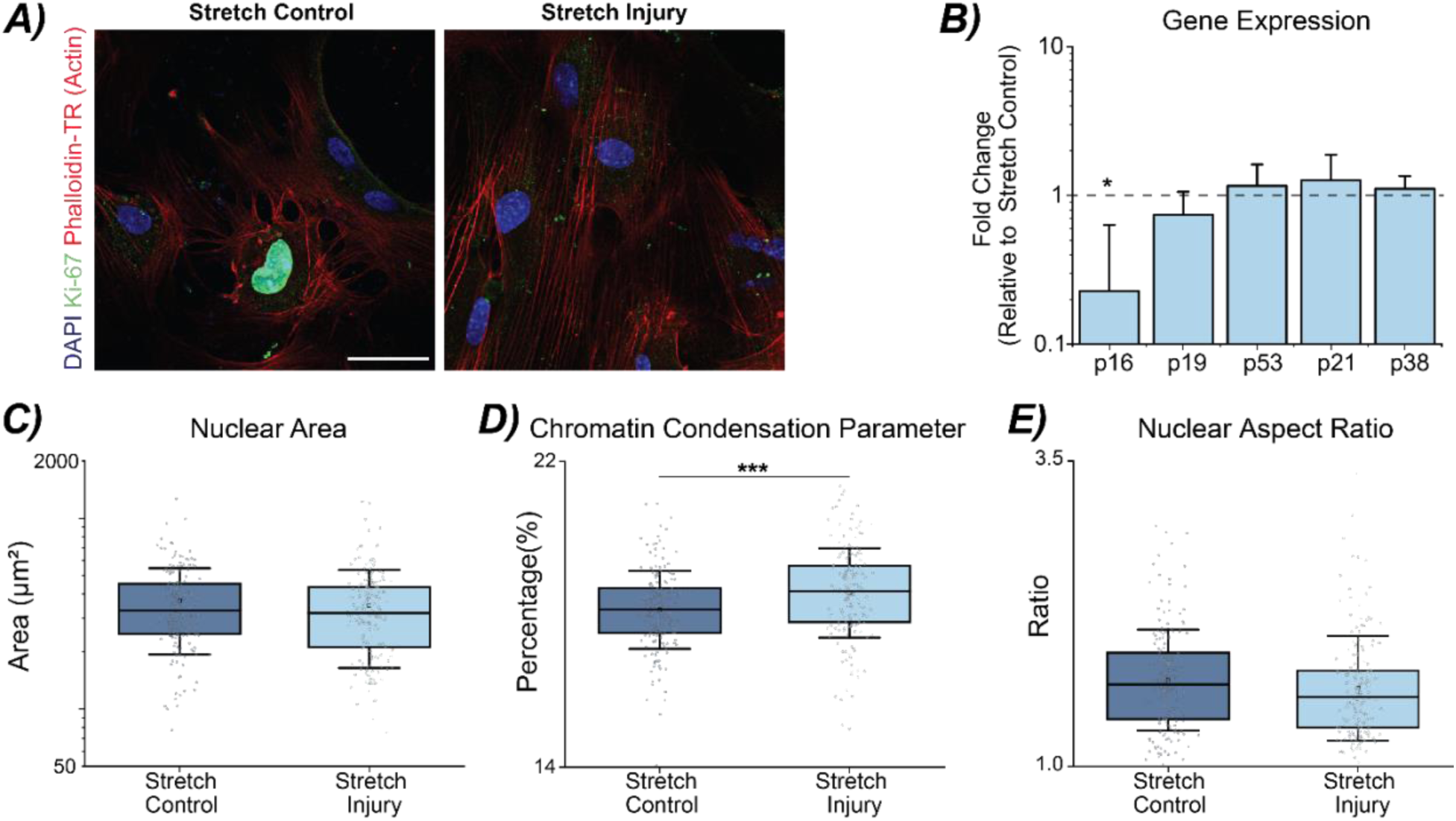
Morphological features of lamin A/C null CFs subjected to Stretch Control and Stretch Injury regimes. (A) Representative area images of lamin A/C null CFs in Stretch Control and Stretch Injury cultures stained with DAPI (405nm), Ki-67 (488nm), and Phalloidin Texas-Red (Actin, 561nm). N=5 animals, Scale bar = 50µm. (B) Fold change of Stretch Injury lamin A/C null CFs to Stretch Control lamin A/C null CFs. A significant downregulation of p16 was found between the two stretch regimes. N = 6 animals. Error bar = SEM. Linear mixed model, ANOVA. *p<0.05. (C). No significant difference is found in nuclear area between Stretch Control and Stretch Injury lamin A/C nullCFs. Plot shows log axis of nuclear area (same data as Fig. 5B). N = 7 animals, n ≥ 25 nuclei/treatment. Error bar = 1 Std. (D) A significant increase in chromatin condensation parameter is observed in Stretch Injury lamin A/C null CFs. N = 7 animals, n ≥ 25 nuclei/treatment. Error bar = 1 Std. ***p< 0.001. (E) Nuclear aspect ratio does not change with stretch regime for lamin A/C null CFs. N = 7 animals, n ≥ 25 nuclei/treatment. Error bar = 1 Std.

**Fig. S4.**
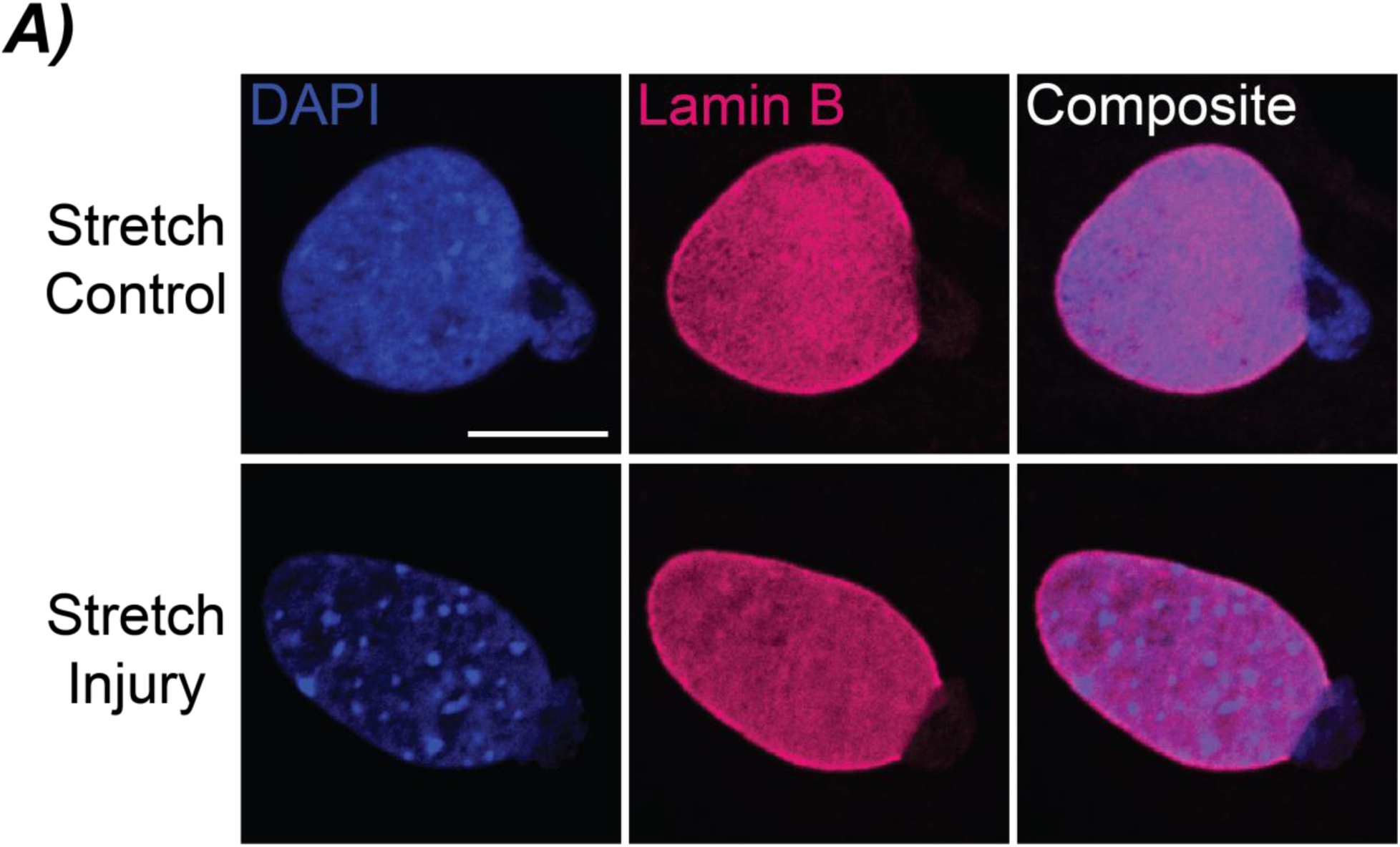
Nuclear rupture observed in mechanical stretched lamin A/C null CFs. (A) Representative images of Stretch Control and Stretch Injury lamin A/C null CFs. DAPI-stained nuclei show nuclear content expelling from the nucleus which corresponds to the same region of lamin B dilution along the nuclear envelope. Scale bar = 10µm.

**Fig. S5.**
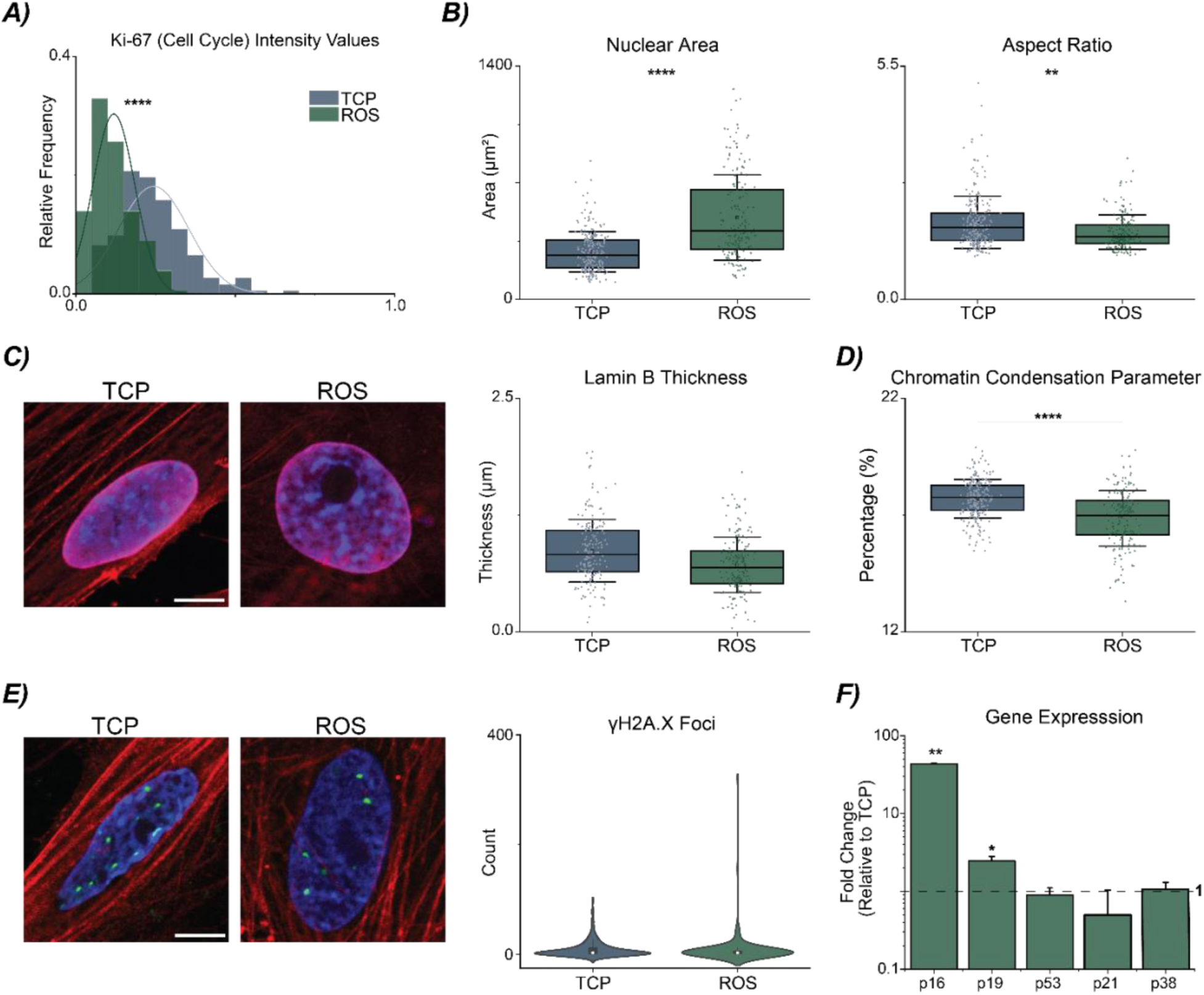
ROS-treated lamin A/C null CFs display several markers of senescence except changes in lamin B thickness and γH2A.X foci counts. (A) ROS-induced lamin A/C null CFs have significant decreased in Ki-67 positive cells compared to TCP CFs. N = 7 animals/group, n ≥ 25 nuclei/treatment. (B) Significant alterations in nuclear area and aspect ratio observed between TCP and ROS-induced lamin A/C null CFs have increased nuclear area and a decrease in aspect ratio. N = 7-9 animals/group, n ≥ 25 nuclei/treatment, Error bar = 1 Std. (C) Representative images of lamin A/C null CFs stained with DAPI (405nm), Phalloidin Texas-Red (Actin, 561nm), and lamin B (640nm). Quantifying the lamin B ring thickness around the nucleus showed no significant difference between TCP and ROS-induced lamin A/C null CFs. N = 7 animals/group, n ≥ 25 nuclei/treatment, Error bar = 1 Std. Scale bar = 10µm. (D) ROS-induced lamin A/C null CFs had decreased chromatin condensation compared to TCP CFs. N = 7-9 animals/group, n ≥ 25 nuclei/treatment, Error bar = 1 Std. (E) Representative images show ɣH2A.X (488nm) staining in TCP and ROS-induced lamin A/C null CFs. Similar to WT ROS-induced senescent CFs, no significance was observed between lamin A/C null CFs in TCP and ROS-treated groups. N = 7-8 animals/group, n ≥ 25 nuclei/treatment, Error bar = 1 Std. Scale bar = 10µm. (F) The gene expression profile of the ROS-induced lamin A/C null CFs has significantly increased p16 and p19 expression. N = 6 animals. Error bar = SEM. Linear mixed model, ANOVA. ****p <0.0001, **p<0.01, *p<0.05.

**Fig. S6.**
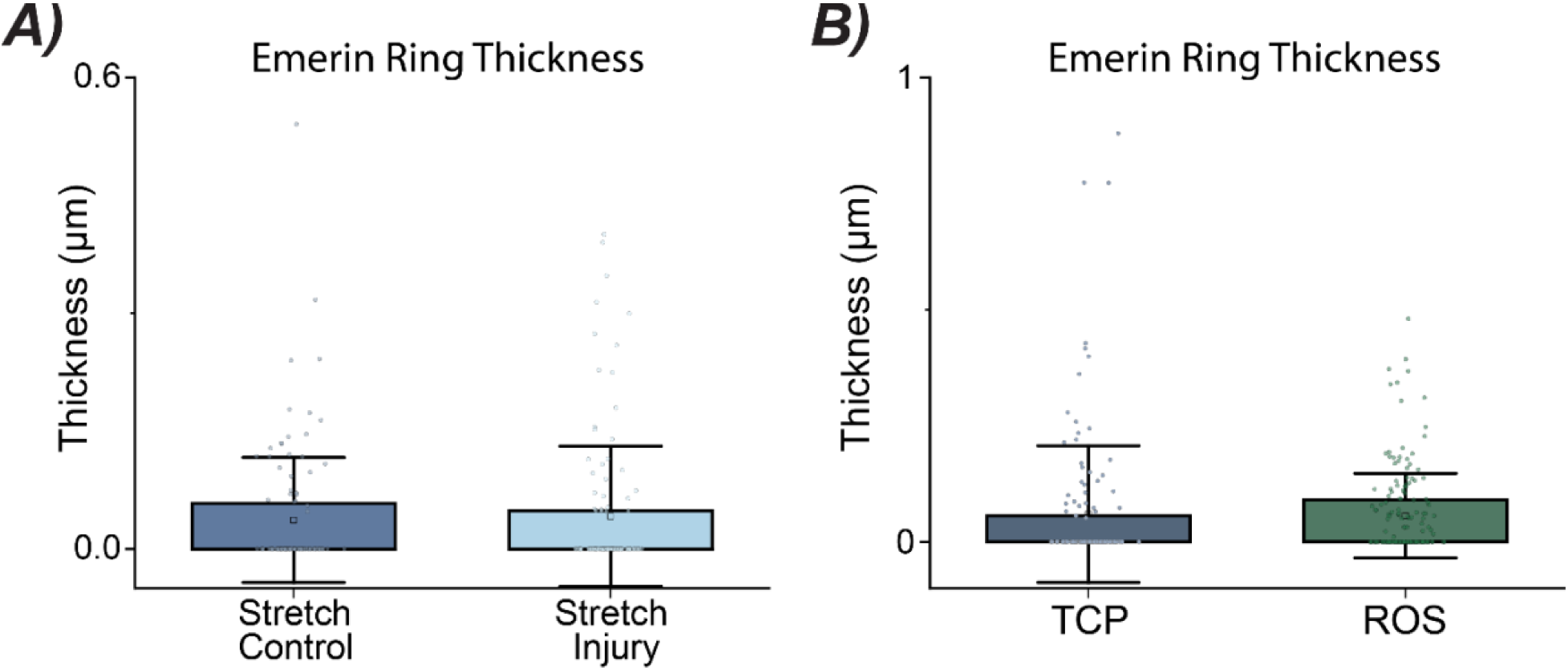
Emerin differences not observed in stretched and ROS-induced lamin A/C null CFs. (A) Quantified emerin ring thickness was similar in Stretch Control and Stretch Injury lamin A/C null CFs. N = 5 animals. n ≥ 25 nuclei/treatment, Error bar = 1 Std. (B) No difference in quantified emerin ring thickness was observed in lamin A/C null CFs on TCP or treated with hydrogen peroxide. N = 5-7 animals, n ≥ 25 nuclei/treatment, Error bar = 1 Std.

**Table S1:**
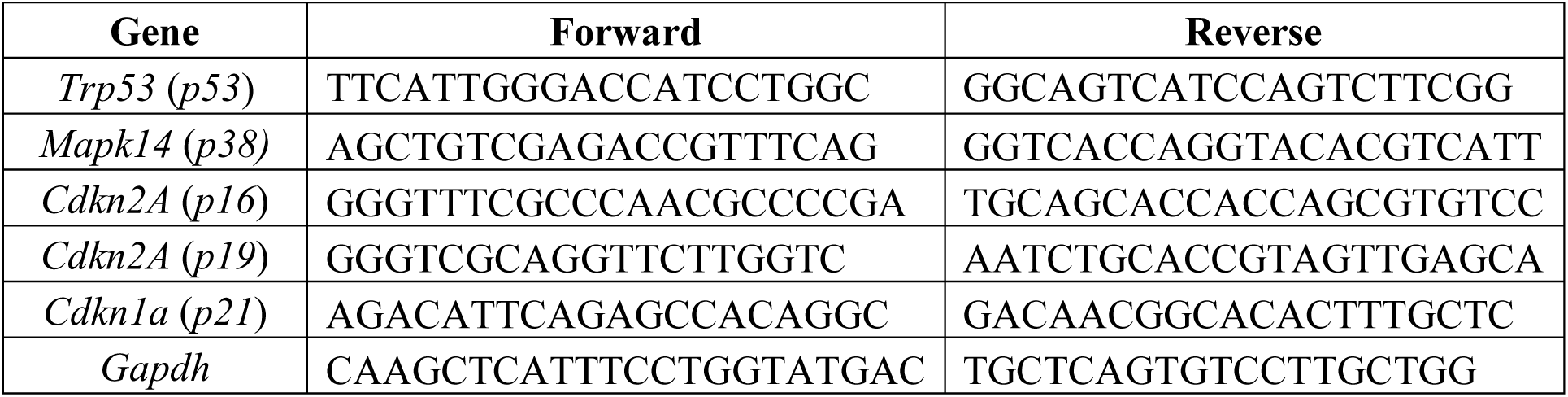
Primer sequences for qPCR of selected genes.

